# On a solution of the biodiversity paradox and a competitive coexistence principle

**DOI:** 10.1101/003095

**Authors:** Lev V. Kalmykov, Vyacheslav L. Kalmykov

**Author notes:** Corresponding author. E-mail address (V. L. Kalmykov).

## Abstract

The biodiversity paradox is the central problem in theoretical ecology. The paradox consists in the contradiction between the competitive exclusion principle and the observed biodiversity. This contradiction is the key subject of the long-standing and continuing biodiversity debates. The paradox impedes our insights into biodiversity conservation. Previously we proved that due to a soliton-like behaviour of population waves complete competitors can indefinitely coexist in one closed homogeneous habitat on one and the same limiting resource under constant conditions of environment, without any trade-offs and cooperations. As this fact violates the known formulations of the competitive exclusion principle we have reformulated the principle. Here we explain why this reformulation of the principle results in a solution of the biodiversity paradox. In addition, we generalize the competitive exclusion principle. Reasoning by contradiction, we formulate a generalized principle of competitive coexistence. These principles expand theoretical basis for biodiversity conservation and sustainable development.

---

There is a long-standing question in theoretical ecology – How so many superficially similar species coexist together? [1, 2, 3] Trying to answer on this question ecologists came to the biodiversity paradox [2, 4, 5]. In theory, according the competitive exclusion principle, complete competitors cannot coexist [6]. In practice there are examples of such coexistence: rain forests [7], coral reefs [8], grasslands [9], plankton communities [2, 4]. The paradox consists in the contradiction between the competitive exclusion principle and the observed biodiversity. “The apparent contradiction between competitive exclusion and the species richness found in nature has been a long-standing enigma” [2]. “Resolving the diversity paradox became a central issue in theoretical ecology” [5]. The key question in this long-standing problem is a validity of the competitive exclusion principle. “Is it true?” [6]

With the help of our deterministic cellular automata models of interspecific competition we strictly logically verified the most known definitions of the competitive exclusion principle. Our results revealed a violation of these definitions and we were forced to reformulate this principle as follows [10]:

> *If each and every individual of a less fit species in any attempt to use any limiting resource always has a direct conflict of interest with an individual of a most fittest species and always loses, then, all other things being equal for all individuals of the competing species, these species cannot coexist indefinitely and the less fit species will be excluded from the habitat in the long run*.
>
> — (*Definition 1*)

*Interspecific resource competition* is a competition between individuals of different species for the same limiting resources in one ecosystem. Under the *limiting resource* we mean a *scarce essential resource*, which directly limits a reproduction of individuals of the species. The *essential resource* is a resource that is fundamentally necessary for existence, functioning and reproduction of individuals of the species and that cannot be replaced by another resource of the species’s intrinsic ecosystem.

According to the Definition 1 an implementation of competitive exclusion of a species in nature requires too strict conditions. Implementation of the competitive exclusion of a species is especially improbable in tropical rainforests as high incoming solar radiation and water availability are the likely drivers of biodiversity. The stringency of conditions of competitive exclusion in according to the Def. 1 eliminates the contradictions between the principle and the observed biodiversity. This is a solution of the biodiversity paradox. The solution of the paradox became possible owing to creation of the strictly mechanistic models of resource competition and reformulation of the competitive exclusion principle [10].

Here trying to simplify, generalize and unify the mechanistic Def. 1 we formulate the following definition of the competitive exclusion principle for any number of resource competitors of any nature:

> *If a competitor completely blocks any use of at least one essential resource by all its competitors, then, all other things being equal, this competitor will exclude them all*.
>
> — (*Definition 2*)

This generalized form of the competitive exclusion principle allows us to interpret different mechanisms of coexistence in terms of availability of access to an essential resource. The both formulations (Def. 1, 2) help to understand a threat to biodiversity that may arise if one competitor will control use of an essential resource of an ecosystem. Today humankind is becoming such a global competitor for all living things [11, 12, 13].

To expand the theoretical basis for biodiversity conservation we formulate here a principle of competitive coexistence. A theoretical fact of indefinite coexistence of complete competitors on the basis of soliton-like interpenetration of population waves [10] demonstrates a possibility of formulation of such principle. This ‘soliton-like’ mechanism implements well-timed regeneration of all essential resources and their optimal allocation among competitors. It enables competitors to avoid direct conflicts of interest. Starting from this mechanism of coexistence of complete competitors and reasoning by contradiction regarding the generalized competitive exclusion principle (Def. 2), we formulate here the second generalized principle – a principle of competitive coexistence, which is also valid for any number of resource competitors of any nature:

> *If all competitors have access to all essential resources and can use them, then, all other things being equal, they will coexist indefinitely or all will perish*.
>
> — (*Definition 3*)

Extinction of all competitors is possible in the case of mutual equal destroying actions or external destructive impact. Indefinite competitive coexistence implies timely access of all competitors to all essential resources. Such access is the necessary condition for biodiversity conservation, sustainable development and existence of any form of life at all.

Previously, we proved that complete competitors can indefinitely coexist in one closed homogeneous habitat on one and the same limiting resource under constant conditions of environment without any trade-offs and cooperations [10]. Coexistence of the competitors in such conditions is possible due to the mechanism of coexistence which is based on the soliton-like behaviour of population waves of competing species. The peculiarities of this mechanism served us as a starting point both for reformulation of the competitive exclusion principle (Def. 1, 2), and for formulation of the principle of competitive coexistence (Def. 3).

The formulated here generalized principles of resource competition expand the theoretical basis for biodiversity conservation and sustainable development. In particular, ecological engineering can be considered as designing of sustainable ecosystems that realize optimal use of all essential resources.

## Acknowledgements

This research was partially supported by an award from the Charity fund of rendering of assistance to scientists “NEW IDEA”.

